# Licensing and competition of stem cells at the niche combine to regulate tissue maintenance

**DOI:** 10.1101/2024.02.15.580493

**Authors:** Rodrigo García-Tejera, Marc Amoyel, Ramon Grima, Linus Schumacher

## Abstract

To maintain and regenerate adult tissues after injury, the numbers, proliferation, and differentiation rates of tissue-resident stem cells must be precisely regulated. The regulatory strategies preventing exhaustion or overgrowth of the stem cell pool, whether there is coordination between different mechanisms, and how to detect them from snapshots of the cell populations, remains unresolved. Recent findings in the Drosophila testes show that prior to differentiation, somatic stem cells transition to a state that licenses them to differentiate upon receiving a commitment signal, but remain capable of fully regaining stem cell function. Here, we build stochastic mathematical models for the somatic stem cell population to investigate how licensing contributes to homeostasis and the variability of stem cell numbers. We find that licensing alone is sufficient regulation to maintain a stable homeostatic state and prevent stem cell extinction. Comparison with previous experimental data argues for the likely presence of regulation through competition for niche access. We show that competition for niche access contributes to the reduction of the variability of stem cell numbers but does not prevent extinction. Our results suggest that a combination of both regulation strategies, licensing and competition for niche access, is needed to reduce variability and prevent extinction simultaneously.

## I. INTRODUCTION

In most tissues, formation, maintenance and regeneration rely on a hierarchical structure in which stem cells, at the apex of the lineage tree, produce differentiating offspring [1, 2]. To do so, a multitude of plausible strategies to regulate their population size have been proposed, such as competition for niche access [3–6], competition for niche signals [6–9], mechanical feedback [10–12], or feedback from the more differentiated populations [13–15]. However, it is still unclear how to distinguish the presence or absence of different regulation strategies from snapshots of the stem cell populations or clonal sub-populations, which are what can often be measured experimentally. Recent years have seen an increasing number of studies reporting a diverse set of internal states in stem cell populations of different tissues [16–25]. The role of cell state heterogeneity in the regulation of stem cell populations and tissue function remains poorly understood.

A startling example where different internal stem cell states are present is spermatogenesis in Drosophila. Spermatogenesis is supported by germline (GSCs) and cyst (CySCs) stem cells [16, 17, 26]. Both CySCs and GSCs coexist in the same niche; a hub consisting of approximately 10 to 12 niche cells providing pro-self-renewal signals [26] (Fig. 1 A). GSCs divide into, in most cases, one daughter GSC that remains in contact with the hub and a differentiating progeny. The latter, known as gonialblast, undergoes four rounds of incomplete division to form a cyst, which later matures into spermatocytes. CySCs, on the other hand, produce post-mitotic differentiating cells, known as cyst cells. Cyst formation begins with a pair of CySCs enveloping a gonialblast, allowing it to progress through differentiation. In turn, the two CySCs differentiate in post-mitotic cyst cells. The regulation mechanisms that promote cyst formation without exhaustion or over-accumulation of GSCs or CySCs remain elusive.

**FIG. 1.**
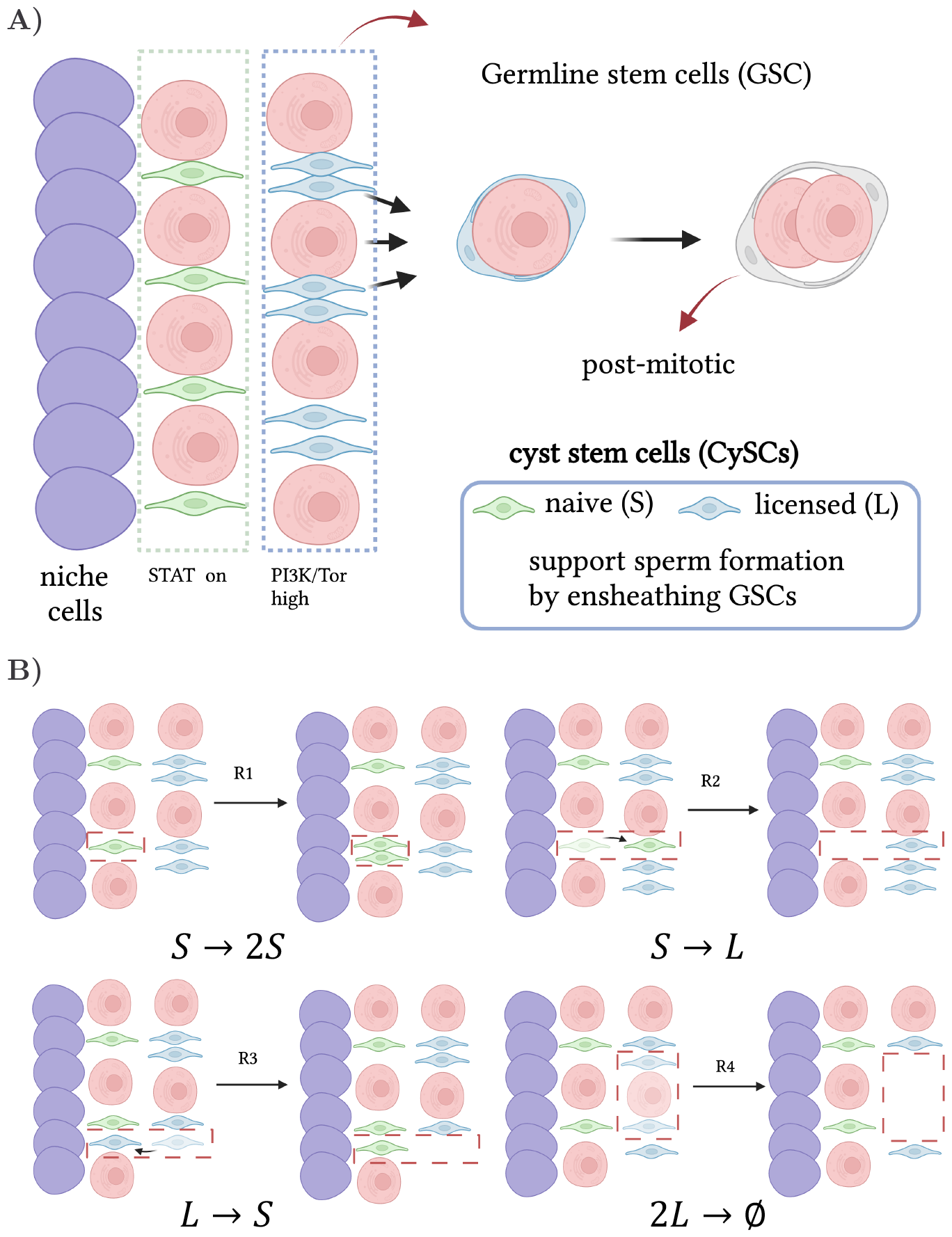
Modelling CySCs in the Drosophila testis. **A)** Schematic of the GSC and CySC niche. GSCs (pink) and CySCs (green) touching the hub (blue cells) get access to pro-self-renewing signals and maintain stemness. CySCs not in contact with the hub lose access to pro self-renewing signals and become licensed for differentiation, although they require active signalling to differentiate. To form a cyst, two CySC daughters surround one GSC daughter (or gonialblast). These co-differentiate, where the CySC daughters become post-mitotic cyst cells, while the germ cell progresses through spermatogonial divisions. **B)** Schematic of the four reactions considered in the *SL* model.

Evidence supports that cyst formation is not regulated by a coordination of GSC and CySC division, but rather at the level of co-differentiation of gonialblasts and CySC daughters [27]. CySCs in contact with the hub maintain self-renewal ability by receiving signals from the hub through the Janus Kinase and Signal Transducer and Activator of Transcription (JAK/STAT) pathway, leading to expression of the transcription factor Zn finger homeodomain 1(Zfh1), which is necessary and sufficient for self-renewal [28]. During differentiation of CySC daughters, Zfh1 expression is downregulated and the cyst cell marker Eyes Absent (Eya) is upregulated [29]. Such commutation of expression levels is triggered by signalling through the Phosphoinositide 3-kinase (PI3K)/Target of Rapamycin (Tor) pathway, which is high for CySCs that are not in contact with the hub [16]. Indeed, knocking down PI3K or Tor pathways results in cells remaining Zfh-1^+^ Eya^−^, confirming that differentiation requires active signalling [17, 30]. Put together, these findings suggest the existence of at least two subpopulations of CySCs, namely a) CySCs in contact with the hub, *naive CySCs*, characterised by high expression of Zfh1 triggered by signalling through the JAK/STAT pathway, and b) CySCs further away from the hub, *licensed CySCs*, with low JAK/STAT activity but high expression of Zfh1. Stem cells in this latter state are ready for differentiation upon receiving an additional, unknown signal through the PI3K/Tor pathway, but can also fully regain stem cell function should they get back in contact with the hub [16, 17].

Whilst the presence of licensed states is thought to play a role in tissue maintenance and regeneration, it is not clear to what extent licensing is a viable regulation strategy for the stem cell numbers, i.e., whether licensing alone is sufficient to provide the homeostatic state with stability and allow the population to recover after injury, or whether additional regulatory mechanisms are needed. To investigate this, we build a minimal mathematical model (*SL* model) in which stem cells can stochastically proliferate, switch back and forth from naive to licensed stem cell states, and form cysts. Analysis of the *SL* model reveals that licensing is a viable strategy to achieve a stable homeostatic state in stem cell populations. We compare our model results with experimental observations of the number of licensed and naive stem cells in adult Drosophila testes under homeostatic conditions. Our analysis shows that the size of the fluctuations for the experimental measurements is lower than those predicted by our model, pointing to the likely presence of additional regulation strategies. We build an additional model (*vSL* model) that includes both regulation through licensing and autonomous regulation of the naive population in the form of competition for niche access, as it has been shown to take place in the CySC/GSC niche [30–32]. The size of fluctuations in the *vSL* model can match the fluctuations size observed experimentally. We show that, while regulation through licensing alone fails at reducing the fluctuations in the size of the stem cell population, it is extremely effective at preventing the extinction of the stem cell pool. Conversely, regulation through competition for niche access alone succeeds at reducing the fluctuations, but cannot prevent extinction. Our results suggest that, for a tissue to exhibit low stochastic fluctuations of the stem cell numbers while simultaneously preventing stochastic extinction, a combination of regulation through licensing and competition for niche access is likely needed.

## II. RESULTS

### A. Licensing of stem cells is a viable regulation strategy

To analyse the viability of licensing as a regulation strategy we built a computational model for stem cell populations that includes both naive and licensed states. The dynamics of stem cell populations has typically been modelled mathematically as critical birth-death processes [3, 33–35], which assumes a minimal set of hypotheses: stem cells *S* undergo proliferation and differentiation/death stochastically, comprising a set of two stochastic reactions that take place at equal rate, proliferation *S* → 2*S*, and differentiation/death *S* → *∅*. One of the reasons for the success of the critical birth-death process to model stem cell populations stems from its ability to reproduce core observations in clonal dynamics, such as the scaling laws relating the number and sizes of surviving clones over time [4, 34–37]. The critical birth-death process assumes that a homeostatic state is present (the self-renewal and differentiation/death rates are equal), but remains agnostic to the mechanism that makes such steady state possible. Recent studies have proposed modifications to the critical birth-death process that includes different mechanisms to ensure homeostasis and recovery dynamics [3, 6, 38].

Our model for the population of CySCs can be seen as an extension to the critical birth-death process, with the following minimal set of hypotheses: **1)** There are two types of CySCs, those in contact with the hub which are proliferative, *naive* stem cells, and those that lose contact with the hub, have no access to niche signals and are ready to differentiate, *licensed* stem cells. **2)** All naive stem cells have the same proliferative potential and can divide stochastically (*R*1 in Fig. 1 B) and can also move away from the niche stochastically, losing contact with the hub, thus becoming licensed stem cells (*R*2 in Fig. 1 B). **3)** Licensed stem cells, on the other hand, can stochastically move back to the niche and regain contact to the hub, becoming naive stem cells again (*R*3 in Fig. 1 B), or they can act in pairs forming a cyst, thus irreversibly differentiating to a post-mitotic state (*R*4 in Fig. 1 B). **4)** We assume that cell death is much more infrequent than any of the four actions *R*1 − *R*4, and can be disregarded (as supported by [39]). Taken together, these hypotheses can be captured by the *SL* model consisting of four stochastic reactions acting upon the naive (*S* species) and licensed (*L* species) populations (Fig. 1 B): *S* → 2*S* (proliferation), *S* ⇌ *L* (licensing and de-licensing), and 2*L* → *∅* (cyst formation). Note that in the *SL* model cyst formation only depends on the number of licensed stem cells, hence, we are assuming abundance of germline stem cells.

A deterministic analysis of the *SL* model, i.e., ignoring the variability of the stem cell numbers due to randomness of the processes involved, reveals that there is a homeostatic state provided that the rate at which naive stem cells proliferate is lower that the rate at which they become licensed (to avoid accumulation of naive stem cells). The parameters of the deterministic *SL* model can be expressed in terms of the average number of naive and licensed stem cells in homeostasis, 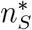 and 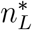 respectively, and the ratio between the average licensing and proliferation rates per cell, *α* (see SI). We can determine the first two parameters from experimental data, obtaining 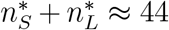 and 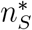 between 13 and 19 (see Methods). For *α*, however, there is no experimental estimate, other than the observation that it must be larger than 1 to avoid accumulation of naive stem cells. We proceed, thus, by studying the behaviour of the *SL* model for plausible values of *α*.

The *SL* model predicts recovery to the homeostatic state after a perturbation regardless of the parameter values (see SI). To analyse qualitatively the mechanisms behind the recovery capabilities we performed in-silico perturbation experiments. In homeostasis, the number of naive CySCs licensing per unit time is balanced with the number of naive CySCs proliferating plus the number of licensed CySCs de-licensing per unit time. If the number of naive CySCs falls below the homeostatic value, the frequency (total number of events per unit time) of both the proliferation and licensing events decreases while the frequency of de-licensing remains constant (Fig. 2 A, left). As a consequence, a de-licensing flux of cells restores the naive CySC pool numbers. In turn, the licensed CySC pool gets partially depleted before converging back to its homeostatic value (Fig. 2 A), right). Conversely, a surplus of the number of naive CyCSs increases the net flux of naive CySCs licensing, followed by a higher increase in the cyst formation rate that restores the numbers of both the naive and licensed CySCs to homeostatic values (Fig. 2 B).

**FIG. 2.**
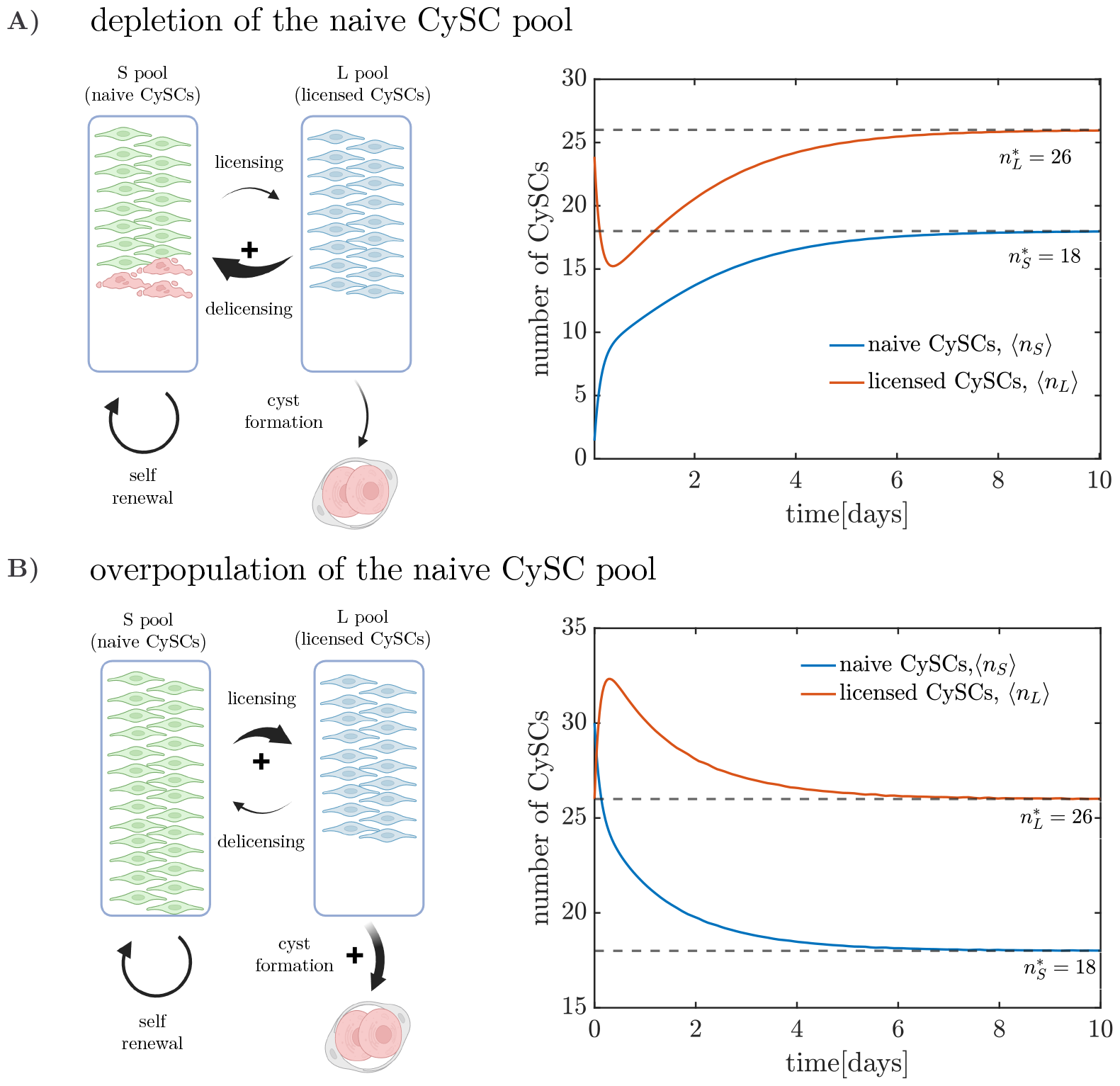
Response of the deterministic *SL* model to perturbations around cell numbers at homeostasis, given by the steady state 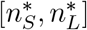. **A)** Upon depletion of the naive CySC pool, a net flux of cells from *L* to *S* state takes place, followed by proliferation restoring back the homeostatic CySC numbers. **B)** When the naive CySC pool is overpopulated a net flux from the *S* to the *L* pool takes place, along with a larger depletion of the *L* pool due to cyst formation. Parameters are set to 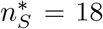, 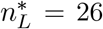 (in accordance with the experimentally observed values) and *α* = 3, with initial conditions [*n*_*S*0_, *n*_*L*0_] = [0, 26] for naive CySC depletion (top) and [*n*_*S*0_, *n*_*L*0_] = [30, 26] for naive CySC overpopulationn (bottom).

The mechanisms of recovery dynamics presented above require the capability of licensed CySCs to regain naive CySC status. In an alternative scenario in which CySCs irreversibly license or differentiate after losing contact with the niche the homeostatic state would be neutrally stable, i.e., the stem cell numbers would not recover after perturbations unless additional regulation strategies are triggered (see SI).

### B. Licensing prevents stochastic extinction and aids recovery dynamics

The stochastic nature of proliferation, licensing, and cyst formation events renders chance extinction of the CySC population plausible. A stochastic implementation of the *SL* model [40, 41] reveals the naive and licensed CySC numbers initially fluctuating about the values of 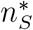 and 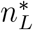 respectively, following a quasi-steady state distribution, but eventually reaching stochastic extinction (Fig. 3 A). Note that extinction is certain to take place for all parameter values and it is not predicted when approaching the *SL* model deterministically, as it is a purely stochastic trait. A more thorough analysis of the onset of extinction dynamics triggered by fluctuations can be found in refs. [3, 42].

**FIG. 3.**
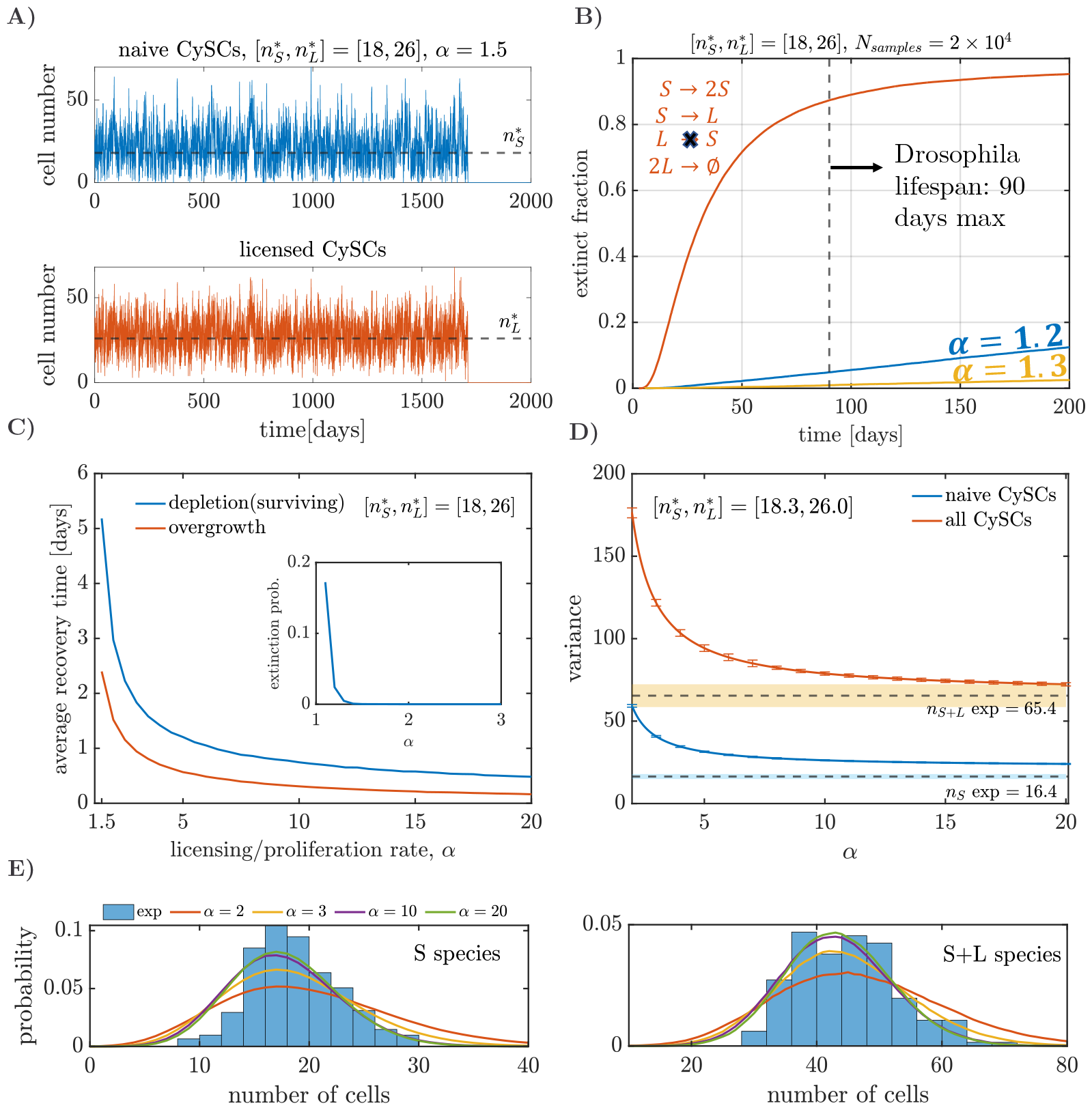
Stochastic implementation of the *SL* model. **A)** Stochastic trajectories for the naive (blue) and licensed (orange) CySCs simulated via the Gillespie algorithm. Cell numbers fluctuate around their homeostatic values (black dashed lines) until they undergo stochastic extinction. **B)** Fraction of extinct tissues as a function of time, over an ensemble of 2 *×* 10^4^ simulated tissues, for the *SL* model (blue and yellow lines) and the modified model scenario (orange lines) in which naive stem cells irreversibly differentiate upon losing contact with the niche (see main text). **C)** Average recovery time (over 2 *×* 10^4^ realisations) after depletion (conditional to survival, blue line) or duplication (orange line) of the naive CySC pool, as a function of *α*. The inset shows the probability of extinction after depletion of the naive CySC pool, as a function of *α*. **D), E)** Comparison between the outcomes of the *SL* model and experimental observations. **D)** Variance of the population sizes (solid lines) predicted by the *SL* model, calculated via Eqs. (22). Error bars indicate standard error. Dashed lines show average experimental measurements, and shaded regions one standard deviation. **E)** Experimental (histogram) and *SL* model (solid lines) distributions of naive (left) and total (right) CySCs numbers.

After establishing that extinction is certain to take place, the question remains whether the expected extinction time falls within the lifespan of Drosophila (median between ∼ 40 and ∼ 90 days [43, 44], although the lifespan is sensitive to temperature and resource availability). For the purpose of this analysis we consider a maximum lifespan of 90 days, although our conclusions remain valid for different values. To investigate the extinction times predicted by our model we simulated, for different values of *α*, an ensemble of 2 *×* 10^4^ tissues. Analysis of the cumulative distribution of extinction times, i.e, the fraction of extinct tissues at different times, reveals that fewer than 5% of the trajectories go extinct within the lifespan of Drosophila, regardless of the value of *α* (blue and yellow lines in Fig. 3 B). For higher values of *α*, the fraction of extinct trajectories within the lifespan of Drosophila decreases. Indeed, when *α >* 1.3 fewer than 0.1% of trajectories go extinct before 90 days. These results remain valid for all the plausible values of 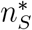 (between 13 and 19).

The effect of licensed CySCs in stochastic extinction becomes evident when considering an alternative scenario where naive stem cells irreversibly differentiate upon loss of contact with the niche. Such scenario can be captured by the reactions *S* → 2*S, S* → *L*, and 2*L* → *∅* (see SI). Simulation of 2 *×* 10^4^ tissues with this model leads to ∼ 90% of the stochastic trajectories going extinct within the lifespan of Drosophila (orange line in Fig. 3 B). The striking difference between the fraction of extinct trajectories in the *SL* model with reversible licensing and lack thereof points to a crucial role of licensing in preventing stochastic extinction.

When a perturbation takes place, the recovery time of the CySC population to its homeostatic state is heavily affected by the licensing and de-licensing rates. To show this, we simulated two types of perturbation experiments, depletion and duplication of the naive CySC pool size. The average recovery time across 2 *×* 10^4^ realisations in both perturbation experiments decreases steadily as *α* increases (Fig. 3 C). Note that for the depletion of the naive CySC pool experiments we consider the recovery time conditional to survival, since there is a chance that extinction occurs before recovery (inset in Fig. 3 C). However, this probability vanishes at very low values (*α* ≈ 1.3). When *α* → 1 the average recovery times increase steadily. Indeed, for the case of depletion of the naive CySC pool, the recovery time diverges. To understand this, note that when the naive CySC pool is empty, the system relies on de-licensing event to repopulate it. Since the ratio between the de-licensing to proliferation rate is 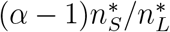 (see SI), letting *α* → 1 while keeping 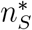 and 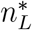 constant leads to the de-licensing rate tending to 0, making the expected time for de-licensing events tend to infinity. The above analysis of the *SL* model illustrates the benefit of having high licensing/de-licensing rates to minimise the probability of extinction after a perturbation (*α >* 1.3 ensures an extinction probability lower than 0.5%) and reduce recovery times.

### C. Licensing alone cannot account for the fluctuations of the stem cell numbers

To explore the effect of licensing and de-licensing in the fluctuations of the stem cell numbers under homeostatic conditions, we compared the fluctuations size from the experimental counts of the number of licensed and naive CySCs (see Sec. IV A) with those from the *SL* model for different values of the ratio between the licensing and proliferation rates of the naive CySCs, *α*. In the *SL* model, when the ratio between the licensing and proliferation rates, *α*, increases while maintaining 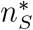 and 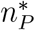 fixed, the variances of the naive 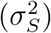 and total 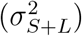 CySC pools decrease steadily (Fig. 3 D and E) approaching the asymptotic values 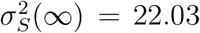 (standard error *SE* = 0.3) and 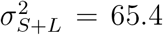 (*SE* = 1.0). For an analytical derivation of the moments of the *SL* model, see SI. These values are larger than the experimentally measured variances 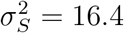 (*SE* = 1.5) and 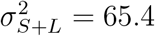 (*SE* = 6.9). In particular, the asymptotic variance of the naive CySC population is 34% higher than its experimental value.

Hence, if the behaviour of the naive and licensed CySC populations is determined solely by stochastic division, licensing/de-licensing, and cyst formation, the observed fluctuations should be larger than the ones observed experimentally. We hypothesise that there are additional regulation strategies or restrictions acting on the CySC populations, which the *SL* model neglects and have been shown to take place in different tissues[3–7].

### D. Introduction of competition for niche access yields fluctuations matching the experimental values

In the case of the Drosophila testes, stem cells are also competing for niche access [30– 32]. In a model that includes competition for niche access, the propensity for stem cell proliferation depends not only on the number of stem cells present, but also on the space available in the niche; a full niche will offer no opportunity for cell growth prior to division, thus rendering proliferation less likely, whilst an empty niche presents no restriction for cell growth and division.

We built upon the *SL* model to include competition for niche access. An efficient way to model stem cell populations with competition for niche access is the birth-death process with volume exclusion (v*BD*) [3]. The vBD includes a proliferation rate that is dependent on the space available by introducing an empty space species *E* and considering that, for proliferation to take place, a stem cell needs to react with an empty space particle, *S* + *E* → 2*S*, an idea introduced in [45]. Hence, when empty space is abundant, proliferation takes place with a rate proportional to the number of stem cells (assuming mass-action kinetics), and when space is scarce the proliferation rate drops, as it has also been explored in Ref. [46]. We can follow similar ideas to introduce competition for niche access in the *SL* model, resulting in the *SL* model with volume exclusion (v*SL*), which consists of three species: naive CySCs *S*, licensed CySCs *L*, and empty space in the niche *E* (Fig. 4). A naive CySC consumes an empty space in the niche to proliferate, *S* + *E* → 2*S*. To license, a naive CySC moves away from the niche, losing contact with niche signals, and leaving an empty space, *S* → *L* + *E*. Conversely, when a licensed CySC goes back to the niche and de-licenses, it consumes an empty space, *L* + *E* → *S*. Similarly to the *SL* model, cyst formation is captured by 2*L* → *∅*. Finally, to account for limited space in the niche, we consider that the number of naive CySCs plus empty spaces is conserved, *n*_*S*_ + *n*_*E*_ = *N*, where *N* is the carrying capacity (maximum number of cells) of the niche. Similarly to the *SL* model, the deterministic behaviour of the v*SL* exhibits a homeostatic state characterised by an equilibrium number of naive 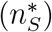 and licensed 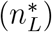 CySCs. The additional parameters are the ratio between licensing to proliferation rates per cell on an empty niche, *α*, and the carrying capacity *N*. In the limit of the carrying capacity *N* → *∞*, the behaviour of the v*SL* and *SL* models become identical. The homeostatic state of the v*SL* model exhibits recovery dynamics after a perturbation (see SI). The recovery dynamics of the v*SL* model involve the cooperation of two mechanisms: licensing and competition for niche access. As a consequence, the recovery time of the v*SL* is lower than in the *SL* model (Fig. 5 A).

**FIG. 4.**
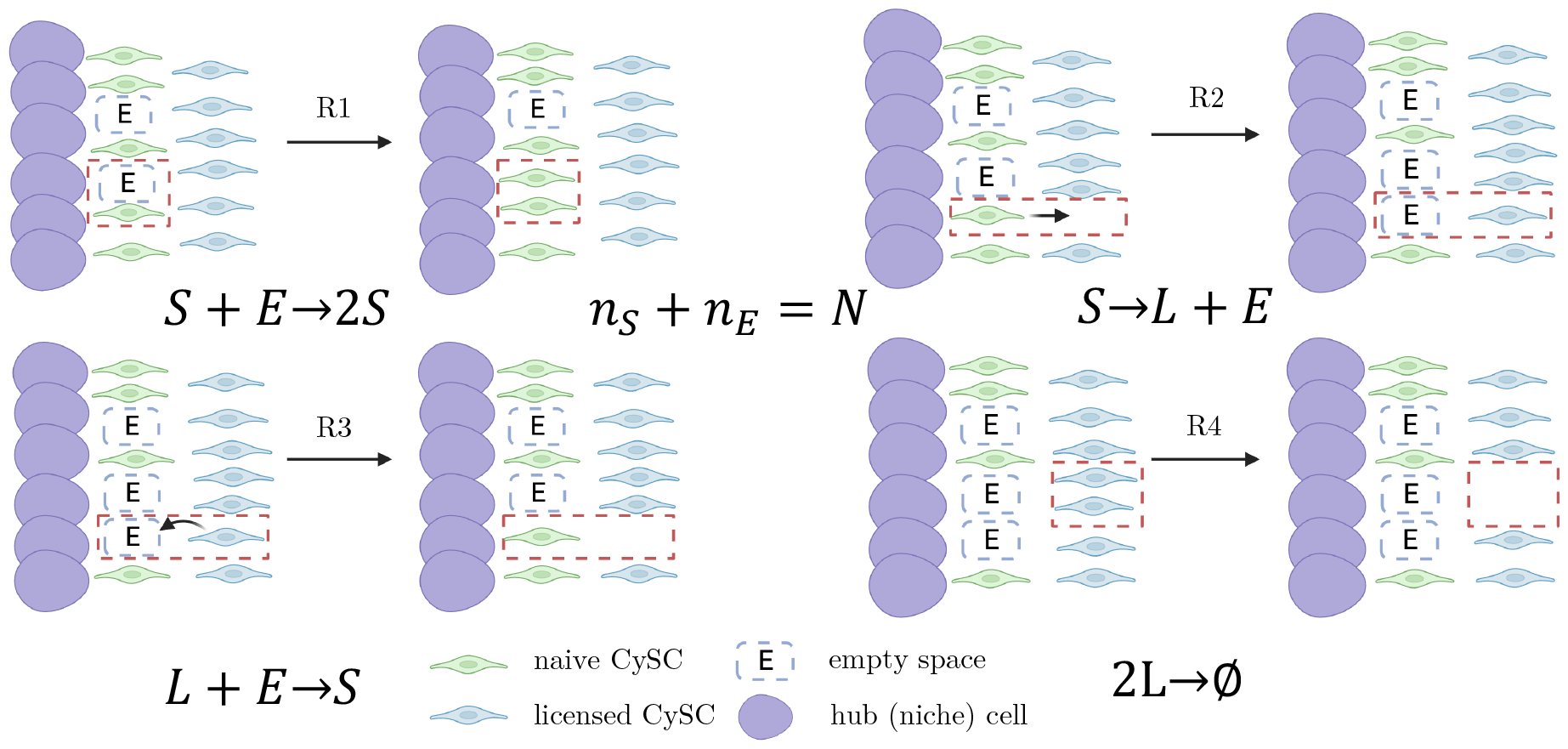
Constitutive actions of the v*SL* model and their corresponding elements in the model’s reaction network. R1 is proliferation, R2 licensing, R3 de-licensing, and R4 cyst formation.

**FIG. 5.**
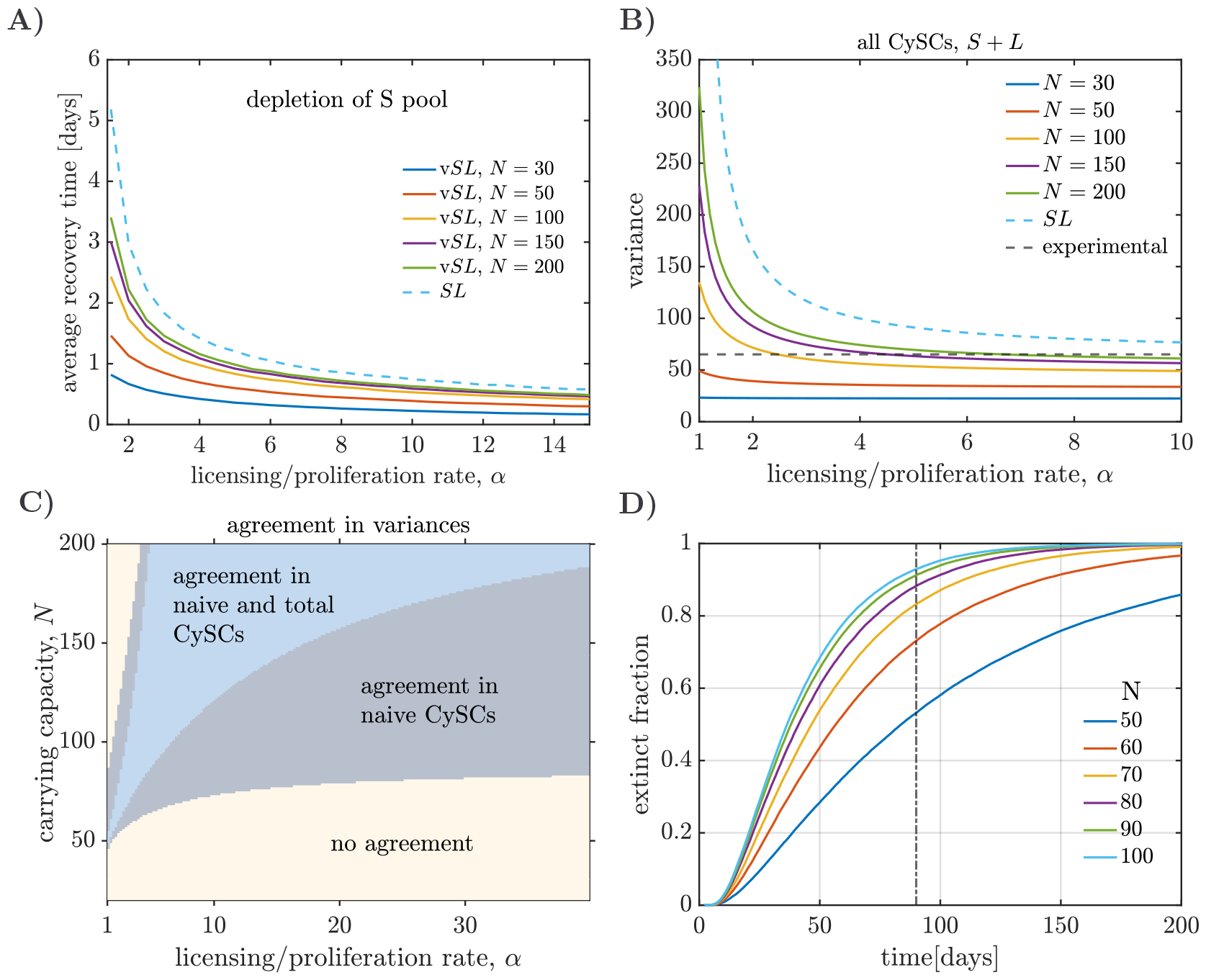
*SL* model with volume exclusion (v*SL*). **A)** Recovery time as a function of the ratio between the licensing and proliferation rates on empty niche, *α*, for the v*SL* model with different carrying capacities (continuous lines). The recovery times are lower than those observed for the *SL* model (dashed blue line). **B)** Homeostatic variance of all (*S* + *L*) CySCs as a function of *α* for different carrying capacities. The variances are always lower than those observed for the *SL* model (blue, dashed lines). Experimentally measured variances (black, dashed lines) fall within the range of variances of the v*SL* model. Similar observations can be made for the variance of the naive CySCs. **C)** Parameter regions of agreement between the experimentally measured variances and the v*SL* model. For parameters in the blue region both the variance of the naive and total CySCs produced by the v*SL* model fall within one standard error of the corresponding variances measured experimentally. The minimum carrying capacity for the parameters that agree with experimental data is larger than *N* = 50. **D)** Extinction experiments for the v*SL* model without the de-licensing reaction but maintaining competition for niche access. The extinct fraction is calculated as the cumulative distribution of extinction times over ensembles of 5 *×* 10^4^ independent simulations.

**FIG. 6.**
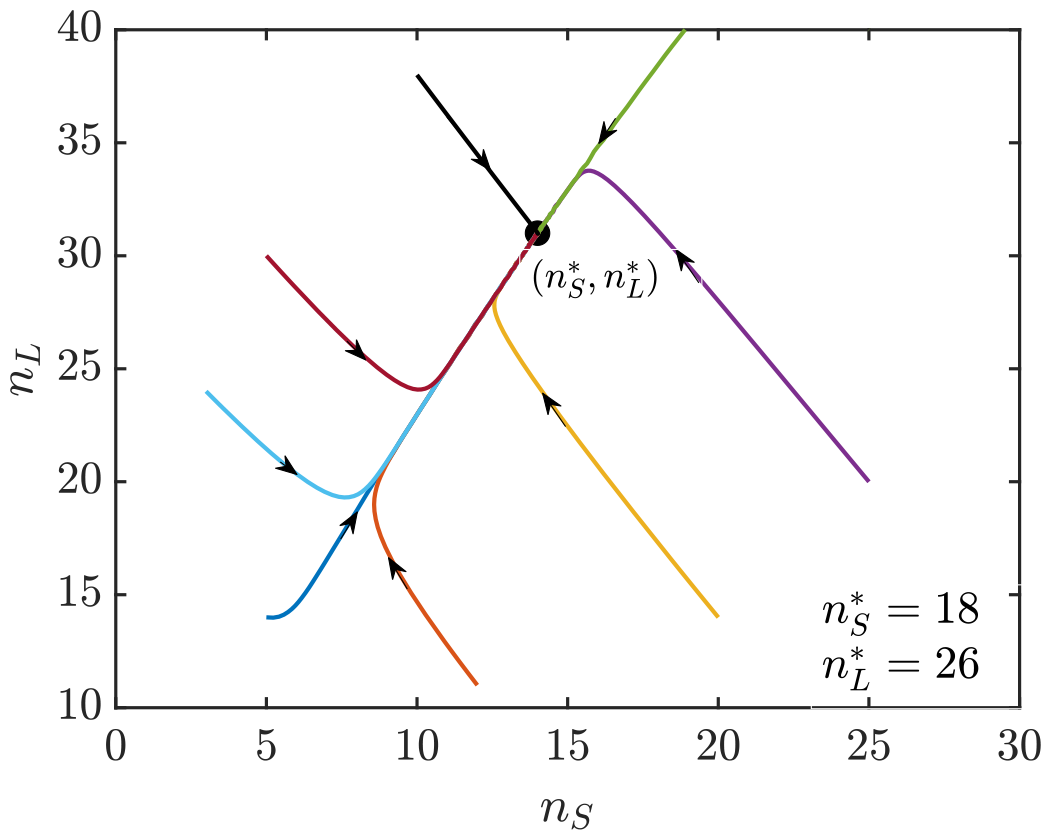
Deterministic implementation of the *SL* model. Trajectories in the *n*_*S*_, *n*_*L*_ configurations space for the mean-field deterministic equations of the *SL* model with 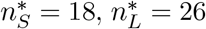 and *α* = 3. Regardless of the initial conditions, the system converges to 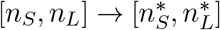.

Combining licensing and competition for niche access as regulation strategies leads to smaller fluctuations of the CySC numbers than solely relying on licensing, as shown by stochastic simulations of the v*SL* model (Fig. 5 B). Analogously to the *SL* model, the fluctuations size is lower for higher values of *α*. Space availability also affects the fluctuations size; rendering them lower for lower carrying capacities *N* (Fig. 5 B). Conversely, in the limit of *N* → *∞*, the distribution of CySC numbers tends to that of the *SL* model.

We found a region of the parameter space, defined by the carrying capacity *N* and licensing/proliferation rates *α*, where the variances for the naive and total CySC population yielded by the v*SL* model agrees with the experimental observations within one standard error (Fig. 5 E). In the same region the variance of the naive CySCs also agrees with experiment. Although our limitations in the sample sizes prevent us from precisely inferring optimal values of *N* and *α*, our analysis establishes a lower boundary for the carrying capacity, which must be larger than 50. A lower carrying capacity leads to fluctuations sizes for the total CySC numbers that are lower than the experimentally observed ones. Note that the carrying capacity serves as an effective parameter in our model, summarising how space availability influences the rate of proliferation per cell. It should not be interpreted as the actual maximum number of cells that can occupy the stem cell niche.

### E. Competition for niche access alone leads to fast extinction

Competition for niche access reduces fluctuations in cell number in a scenario where CySCs can reversibly transition between naive and licensed states. The question remains whether competition for niche access alone is capable of sustaining homeostasis preventing stochastic extinction. To investigate this, we considered an alternative model that includes competition for niche access but assumes that stem cells irreversibly differentiate upon detachment from the niche. Analogously to Sec. II B, we implemented such scenario by removing the de-licensing reaction in the v*SL* model.

For carrying capacities within the region of agreement between the v*SL* model and the experimental data (i.e., *N >* 50, see Fig. 5 C), competition for niche access alone leads to fast extinction (Fig. 5 D). After simulating an ensemble of 5 *×* 10^4^ tissues, the fraction of extinct tissues over time depends on the carrying capacity (Fig. 5 D). However, for all plausible carrying capacities, more than 50% of the tissues go extinct within the lifespan of Drosophila. In contrast, when de-licensing is present but competition for niche access is absent (*SL* model), fewer than 5% of the tissues go extinct for all parameter regimes (see Fig. 3 B). Competition for niche access alone is thus not capable to prevent stochastic extinction.

## III. DISCUSSION

In this work we have investigated the role of cell states in which stem cells are licensed but not yet committed to differentiation in the maintenance of homeostasis, recovery dynamics after injury, and variability of the stem cell pool size due to randomness in stem cell division and cyst formation. We have focused on the somatic stem cells that participate in spermatogenesis in the Drosophila testes as a paradigmatic example. Analysis of a minimal mathematical model that considers CySCs in naive and licensed states (*SL* model) suggests that the ability of CySCs to transition back and forth between naive and licensed states provides the system with a robust homeostatic state and recovery abilities after a perturbation (Sec. II A). The importance of licensed states in recovery after an injury becomes evident when considering a situation in which stem cells irreversibly differentiate upon losing contact with the niche; in such case the system cannot recover after a perturbation. Moreover, comparison between these two scenarios suggests that the ability of CySCs to reversibly switch between naive and licensed states is crucial to prevent stochastic extinction (Sec. II B). However, the variability in the stem cell numbers observed experimentally is significantly lower than what would be expected from stochastic naive CySC division, licensing/de-licensing and cyst formation; thus pointing to the presence of additional regulation (Sec. II C). Incorporating competition for niche access leads to the v*SL* model, which can yield lower fluctuation sizes, matching the experimentally measured values (Sec. II D). Although competition for niche access is effective at reducing the fluctuations size, its presence in a scenario where CySCs irreversible differentiate upon losing contact with the niche leads to stochastic extinction, typically within the lifespan of Drosophila (Sec. II E). A combination of both regulation strategies is needed to reduce the variability of the stem cell numbers due to randomness in the cell division, licensing/de-licensing and cyst formation processes, while preventing stochastic extinction of the CySC population simultaneously.

Minimal mathematical models capture the fundamental behaviours of systems, leaving aside accessory details. Here, we assumed that naive stem cells stochastically proliferate and license, and licensed stem cells stochastically de-license and form cysts, which provided a picture of the behaviour of the stem cell numbers in the absence of any additional mechanism and details. However, both in the *SL* and v*SL* models we assumed that cyst formation depends solely on the CySC population, ignoring the role of GSC availability in the cyst formation rate. This assumption is equivalent to assuming abundance of GSCs, which might not be the case in certain situations. When introducing competition for niche access in the v*SL* model, we considered that naive CySCs compete with each other for access to niche signals, ignoring the presence of naive GSCs also competing. This consideration a mounts to assuming that the naive GSC numbers remain constant through the process. The effect of the GSCs can be incorporated to the v*SL* model through a varying carrying capacity that is reduced after a GSC proliferation event and increased after a licensing one. Our modelling of competition for niche access was inspired in the birth-death process with volume exclusion model [3]. This model treats cells as hard spheres without any ability to deform, while CySCs have certain deformation potential. Future refinement of the v*SL* model could relax the level of volume exclusion, for example by allowing occasional proliferation and de-licensing in the absence of empty spaces. The study of the combined dynamics of CySCs and GSCs, where both cell lines can license and de-license, compete for the same space within the niche, and have different deformation abilities, is a further complexification that is an interesting avenue for future research.

Stem cell populations with licensed states are present in tissues other than Drosophila testis. For example, licensed states have been reported in spermatogenesis in the mouse testis [20, 47], as well as in intestinal epithelium [21], hematopoietic stem and progenitor cells [1, 19, 22], and bulge stem cells [22]. Licensing is also closely related functionally to priming in embryonic stem cells [18, 23–25]. The *SL* and v*SL* models can provide a basis for quantitative analysis in such systems. As shown in Sec. II C, quantifying the variability of the stem cell numbers in homeostasis due to the randomness in the cell division processes can provide information on the regulatory mechanisms maintaining homeostasis. The models presented here can be adapted to represent other tissues with different mechanisms. Depending on the tissue of interest, the cyst formation reaction may be substituted by an alternative differentiation process, or autonomous regulation of the naive stem cells in open niches may be better modelled through competition for mitogens rather than competition for niche access. Our study provides a useful framework to investigate how homeostasis is maintained in tissues where stem cells can adopt heterogeneous states, and explore plausible pathways leading to regulation break down in disease.

## IV. METHODS

### A. Measurement of CySC numbers

CySC numbers and were obtained from the datasets published in the Results section of Ref. [30]. In brief, two methods of counting were used. In the first, all cells expressing Zfh1 but not Eya were counted as CySCs, giving an estimate of 44 total CySCs (including both licensed and naive populations). Based on clonal labelling the proportion being naive stem cells has been estimated as 14 [30]. In the second, a membrane marker was used to determine whether Zfh1-positive cells were in contact with the niche. These were considered to represent the naive CySC population, as being in direct contact with the niche, yielding an estimate of 18 naive stem cells [30]. As these estimates each come with their specific inaccuracies and potential measurement errors, we consider the range of 13 to 19 naive stem cells (taking into account the standard error of measurements) when looking for agreement between model simulations and experiments.

### B. Deterministic modelling of stem cell population dymaics

To model the CySC population we considered a system with two species, naive (*S*) and licensed (*L*) stem cells. In the *SL* model naive stem cells can proliferate and license, and licensed stem cells can de-license and form cysts in pairs. This behaviour is captured by the set of reactions *S* → 2*S, S* ⇌ *L*, and 2*L* → *∅*. To approach the system deterministically we considered mass-action kinetics under well-mixing and dilute gas conditions, and take a mean field approach. If the licensing rate is larger than the proliferation rate, the deterministic system exhibits a non-zero, stable steady state characterised by 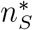 naive and 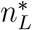 licensed stem cells, and the deterministic equations are given by (see SI)

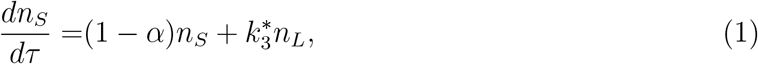

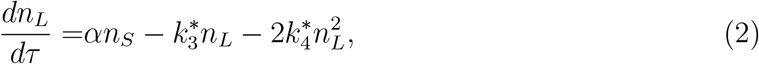

where *τ* is the time rescaled by the proliferation rate and *α* is the ratio between the licensing and proliferation rates.

In the case of the v*SL* model, the defining reactions are *S* + *E* → 2*S, S* ⇌ *L* + *E*, and 2*L* → *∅*, where *E* denotes the number of empty spaces within the niche. To account for limited space in the niche, we considered that the sum of naive stem cells and empty spaces remain constant, *n*_*S*_ + *n*_*E*_ = *N*, where *N* is the carrying capacity of the system. Taking the same assumptions as in the *SL* model, the system has a stable steady state and the deterministic equations are given by (see SI)

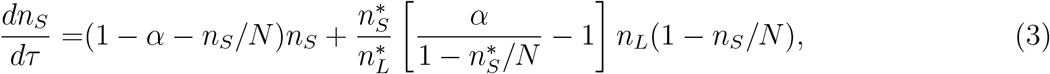

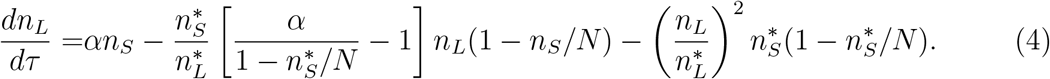

We solve Eqs. (2) and (4) numerically using an adaptive Runge-Kutta (order 4) method.

### C. Stochastic modelling of stem cell numbers

We analysed the stochastic *SL* and v*SL* models by combining simulations and analysis. Stochastic simulations were performed using the Gillespie algorithm [40]. When studying recovery dynamics (Figs. 3 C and 5 B) we defined the recovery time as the time that takes the in-silico tissue to get the homeostatic number of naive CySCs for the first time after the perturbation.

To calculate the variance of the distributions of CySC numbers predicted by the *SL* model we approximately solved the model’s master equations (see SI). Our approximate solution makes use of the linear noise approximation [48–50], and yields analytical expressions for the variance and covariance of the naive and licensed CySC numbers:

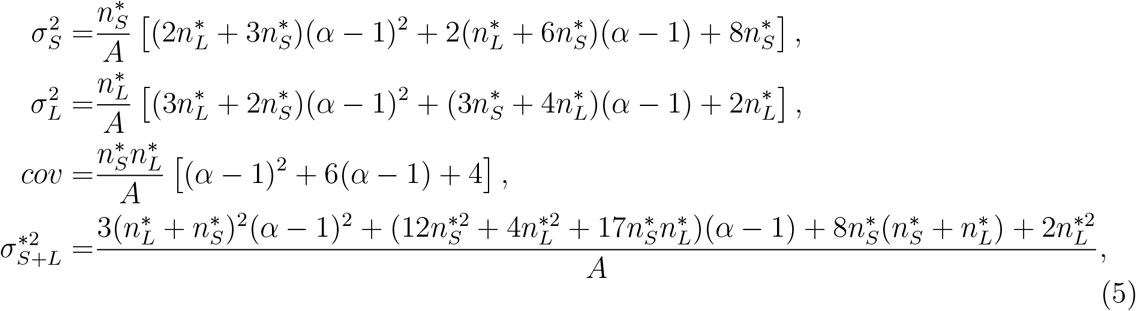

where 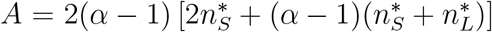. In the case of the v*SL* model the analytical expression for the variances is rather cumbersome. Instead, we performed the linear noise approximation numerically and obtained accurate numerical estimates of the variances of the stem cell numbers according to the v*SL* model (see SI).

## VII. SUPPLEMENTAL INFORMATION

### A. The *SL* model

The *SL* model represents a population of stem cells that can be either in naive (*S*) or licensed (*L*) state. Naive stem cells can proliferate or license stochastically, and licensed stem cells can de-license or act in pairs to form a cyst, leading to the reaction network

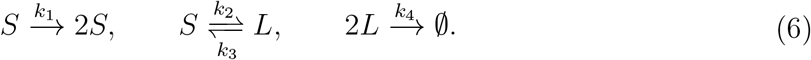

#### 1. Mean-field deterministic equations and stability analysis

To arrive at the deterministic equations of the *SL* model we assume mass-action kinetics, consider the *S* and *L* population to reside in a volume Ω as a dilute and well-mixed gas, and apply a mean-field approximation, leading to the rate equations

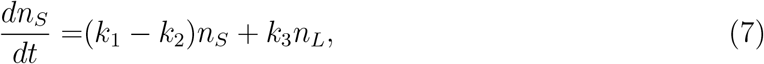

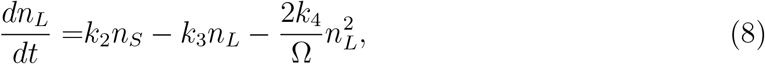

where we assumed the approximation 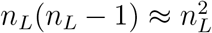. Non-dimensionalizing time *τ* = *k*_1_*t* and defining 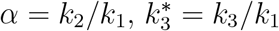 and 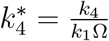 leads to

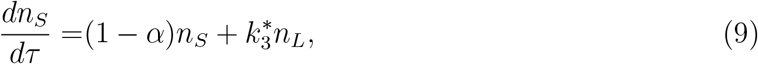

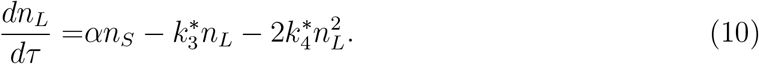

We can see that there is always a trivial steady state 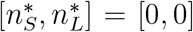. The necessary and sufficient condition for the existence of a nontrivial steady state is that *α >* 1, which means that the proliferation rate must be lower than the licensing rate, to avoid divergence of the *S* population due to over-accumulation. Assuming *α >* 1, the non-trivial steady state is given by

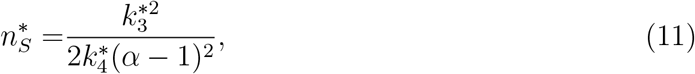

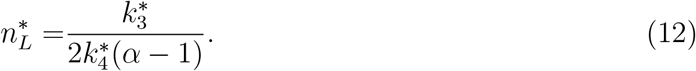

Note that there exists a combination of rate constants that yields any steady state value, 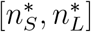. We can re-parameterise our system according to the steady state values, observing that 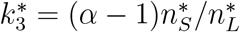 and 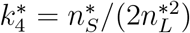, leading to

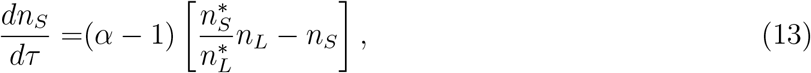

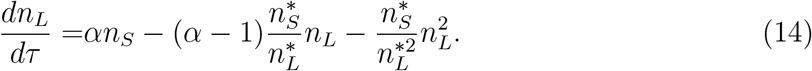

The stability of the steady states can be analysed via the system’s Jacobian

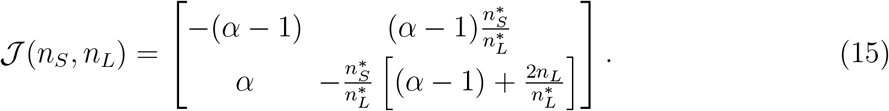

The Jacobian at the trivial steady state, *J* (0, 0), has one positive and one negative eigenvalue, given by

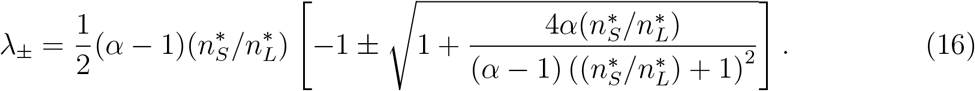

It is simple to prove that the negative eigenvalue is related to perturbations along a direction with negative slope, and the positive one to perturbations along a direction with positive slope. All real perturbations are along directions with slope higher or equal to zero so, in practice, the trivial steady state is repellent. The Jacobian in the non-trivial steady state, 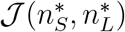, has eigenvalues with negative real parts, given by

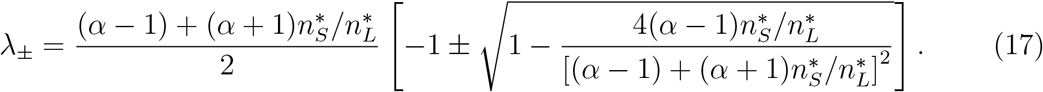

Both eigenvalues are real, since 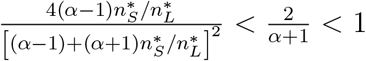. Hence, the nontrivial steady state is always attracting, and exhibits recovery dynamics without oscillations.

#### 2. Stochastic implementation of the SL model

Here, we derive analytical expressions for the second moments and total distribution of the CySC numbers in homeostasis (Eq.(5)). We make use of the linear noise approximation (LNA) to the *SL* model’s master equations [48, 49] and obtain expressions for the second moments of the distribution of CySC numbers in quasi-steady state, which we subsequently use to calculate the parameters of a negative binomial as an approximation for the total distribution of the stem cell numbers.

The Jacobian at the nontrivial steady state of the mean-field deterministic equations is given by evaluating the jacobian *J* given by Eq. 15 at 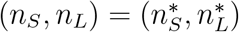:

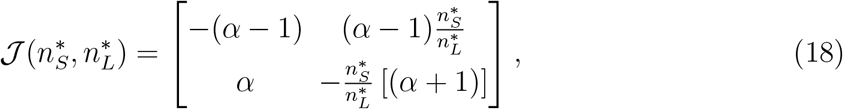

where 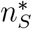 and 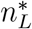 are the homeostatic numbers of naive and licensed CySCs respectively, and *α* is the ratio between the licensing and proliferation rates of naive CySCs. The linear noise approximation amounts to performing the van Kampen’s system-size expansion of the master equation in powers of *N* ^−1*/*2^ and collecting the terms of order *N* ^0^, obtaining a Fokker-Planck equation [48]. For reaction networks consisting of many species (here naive and licensed CySCs), it is convenient to adopt a matrix formulation for the LNA, as introduced in [50]. Under such formulation, the variances and covariances between species in steady state are given by the solution to the Lyapunov equation

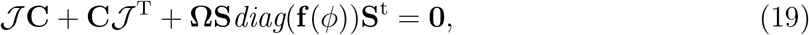

where *J* is the Jacobian of the deterministic equations in steady state, **C** is the variance matrix to be calculated, Ω is the system volume (here taken as Ω = 1 without losing generality), **S** is the stoichiometric matrix and **f** the propensity vector. The stoichiometric matrix of the *SL* model’s reaction network is given by

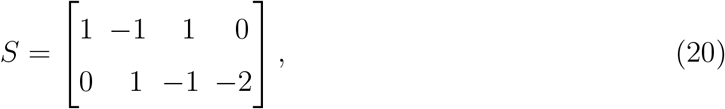

where the first row represents the net changes of *S* species and *L* is encoded in the second row. Considering mass-action kinetics, the propensity vector in steady state is given by

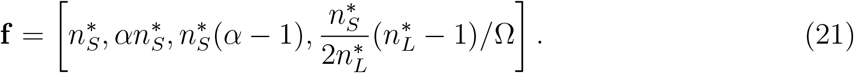

Solving Lyapunov’s equation for the system Jacobian and diffusion matrix yields,

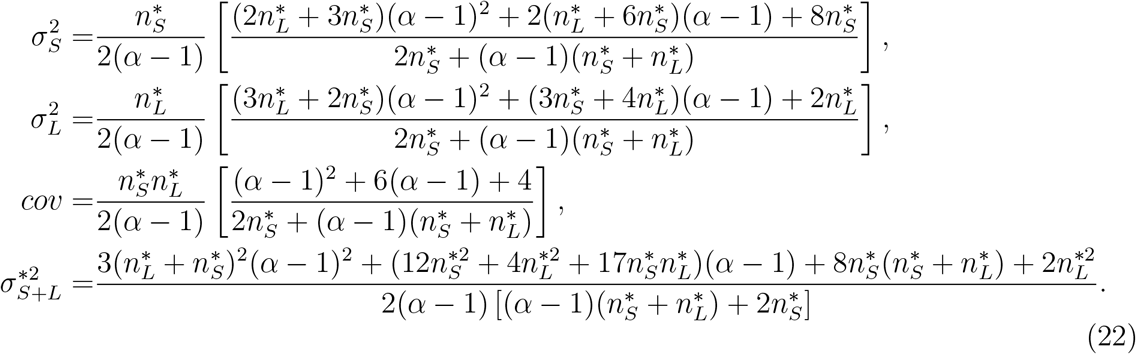

Note that when *α* → 1 the fluctuation sizes tend to *∞*.

Instead of taking the Gaussian approximation yielded by the LNA, we use the expressions for the first two moments of the distributions to calculate the parameters of negative binomials for the probability distributions of the number of naive and licensed CySCs in homeostasis. In contrast to the LNA, a negative binomial is able to capture the skewness of the distributions. The resulting (quasi) steady state distributions for the *S* and *L* species read

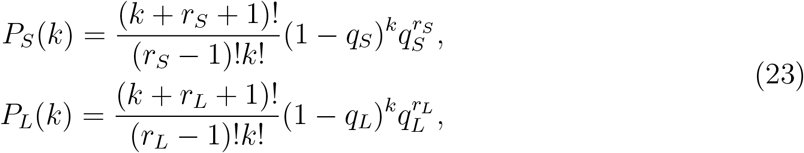

where 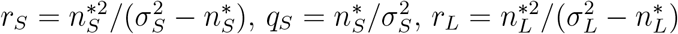 and 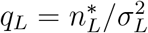. The negative binomial/LNA approximation is in good agreement with stochastic simulations for most values of *α* (Fig. 7 A and B). For lower values of *α* extinction events are more frequent. As a consequence, the probability distribution of the surviving trajectories (captured by the SSA) is biased towards higher stem cell numbers, which creates a mismatch between the first moment of the distributions calculated via the SSA and the NB/LNA approximation (see Fig. 7 B). The second moment, however, remains accurate for most values of *α*, only showing disagreement for *α* ≲ 1.1 (since the LNA prediction diverges when *α* → 1), as shown in Fig. 8).

**FIG. 7.**
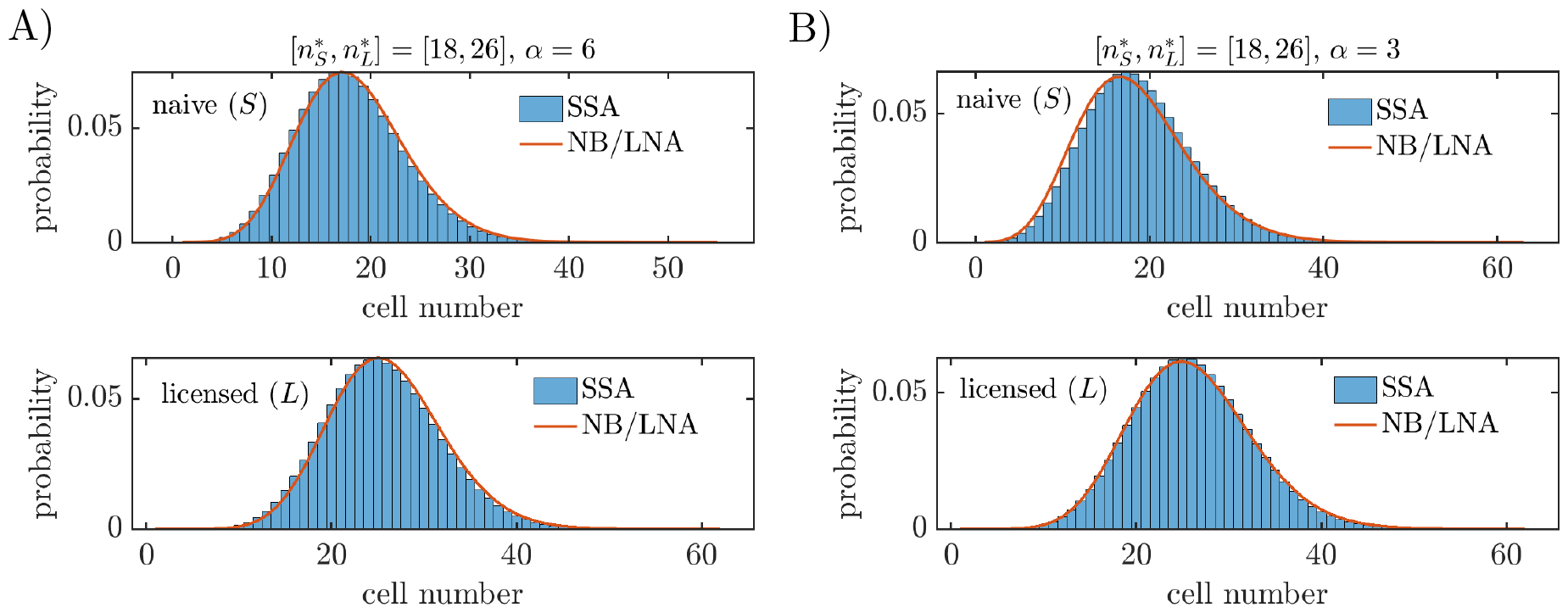
Negative binomial/LNA approximation (red line) to the solution of the Kolmogorov’s forward equation for the *SP* model in steady state (Eq. (23)), for the experimentally estimated values of 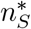 and 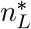, for *α* = 6 (**A**) and *α* = 3 (**B**). Histograms correspond to stochastic simulations (extinction events are excluded).

**FIG. 8.**
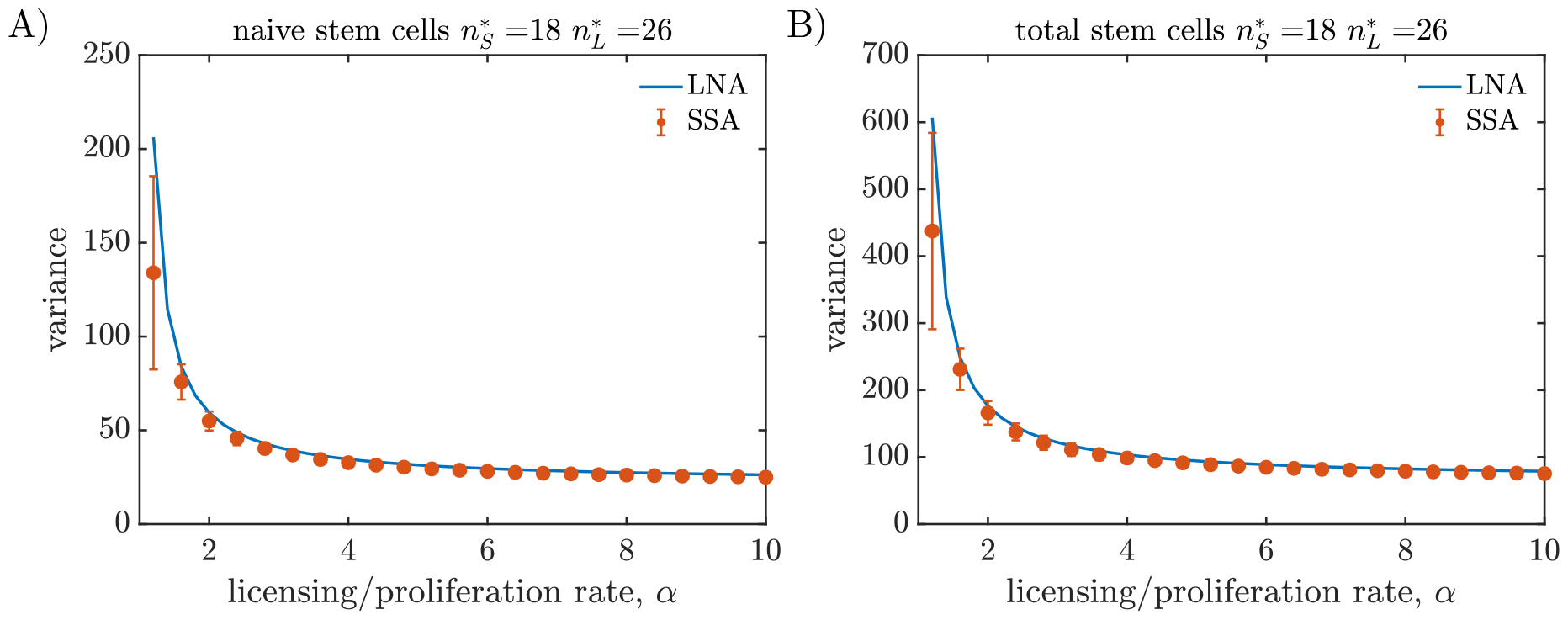
Second moments of the distribution of naive. (**A**) and total (**B**) stem cell numbers in quasi-steady state, as a function of *α* for the *SL* model. The variance measurements (orange dots) are averaged over 5 *×* 10^3^ realisations of the SSA with 1 *×* 10^4^ time points. Error bars show one standard deviation in the ensemble of realisations. The LNA predictions (blue lines), obtained from Eq. (22), are in agreement with the SSA measurements.

### B. Analysis of alternative scenario to the *SL* model

In section II B we analyse an alternative scenario in which naive stem cells irreversibly differentiate when they lose contact with the niche, which can be captured by the reaction network:

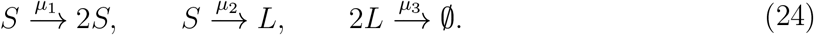

The mean-field deterministic equations of the system are

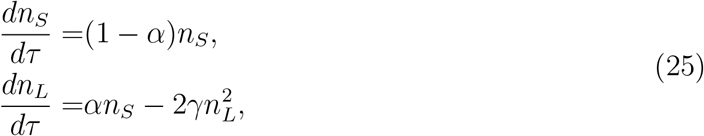

where *α* = *µ*_2_*/µ*_1_, *γ* = *µ*_3_*/*(*µ*_1_Ω), being Ω the system volume, and *τ* = *µ*_1_*t*. For the meanfield equations to have a steady state 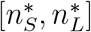 it must be *α* = 1 and 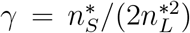. It is simple to prove that the system Jacobian in steady state has eigenvalues *λ*_1_ = 0 and 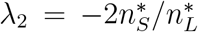, the null eigenvalue implying that the system would not go back to the original steady state after perturbation, as all the points in the curve 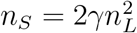 are steady states.

### C. The v*SL* model

The v*SL* model can be seen as an extension of the *SL* model that incorporates competition for niche access in the naive stem cell population, modelled as a volume exclusion effect [3]. Naive stem cells (*S*) can proliferate stochastically, provided that they find empty spaces within the niche (*E*). They can also license by moving away from the niche, thus leaving an empty space available. Licensed stem cells can de-license by taking up an empty space in the niche, or act in pairs to form a cyst, leading to the reaction network

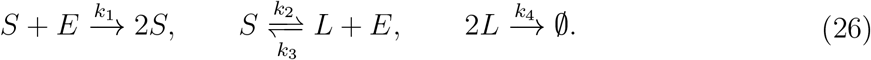

To account for a finite space in the niche, we introduce a carrying capacity *N* by demanding that at all times the number of naive stem cells plus empty spaces remains constant, i.e., *n*_*S*_ + *n*_*E*_ = *N*.

#### 1. Mean-field deterministic equations and stability analysis

Assuming mass-action kinetics, considering the well-mixing and dilute gas hypotheses, and imposing *n*_*E*_ = *N* − *n*_*S*_, where *N* is the niche’s carrying capacity leads to the mean-field equations

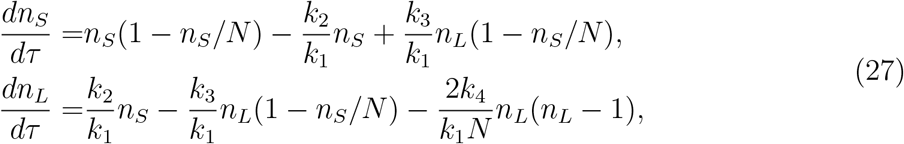

where we have defined the non-dimensional time *τ* = *k*_1_*t*. Let us now assume the existence of a nontrivial steady state characterised by 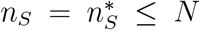 and 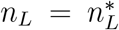, where both 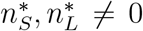. We prove the existence of such a steady state later on. Substitution of the steady state values in the rate equations leads to

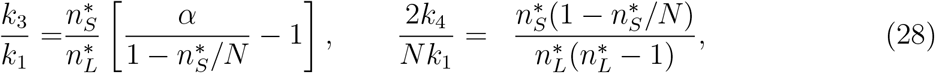

where we have defined *α* = *k*_2_*/k*_1_. Substitution of the identities (28) in the rate equations (27) leads to

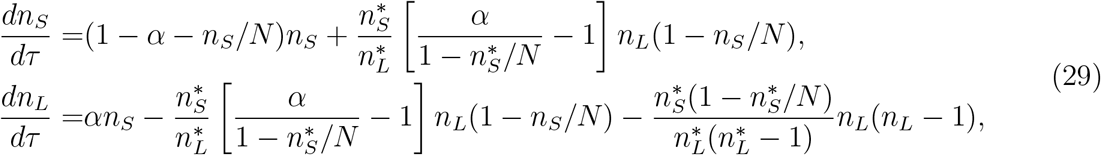

which is Eq. (4) of the main text.

To prove that a nontrivial steady state 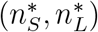 exists consider the nullclines for the rate equations (27), given by

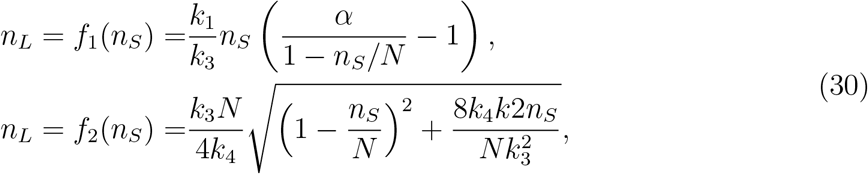

as shown in Fig. 9. The existence of the nontrivial steady state with 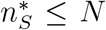 is a consequence of the following conditions, which are fulfilled for *α >* 1: a) *f*_1_(0) = *f*_2_(0) = 0, b) *∂f*_1_*/∂n*_*S*_(0) *>* 0, c) *∂f*_1_*/∂n*_*S*_ → *∞* as *n*_*S*_ → *N* ^−^, 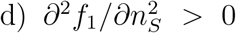 in [0, *N*), 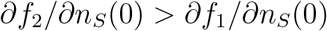, and 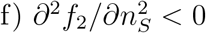 in [0, *N*).

**FIG. 9.**
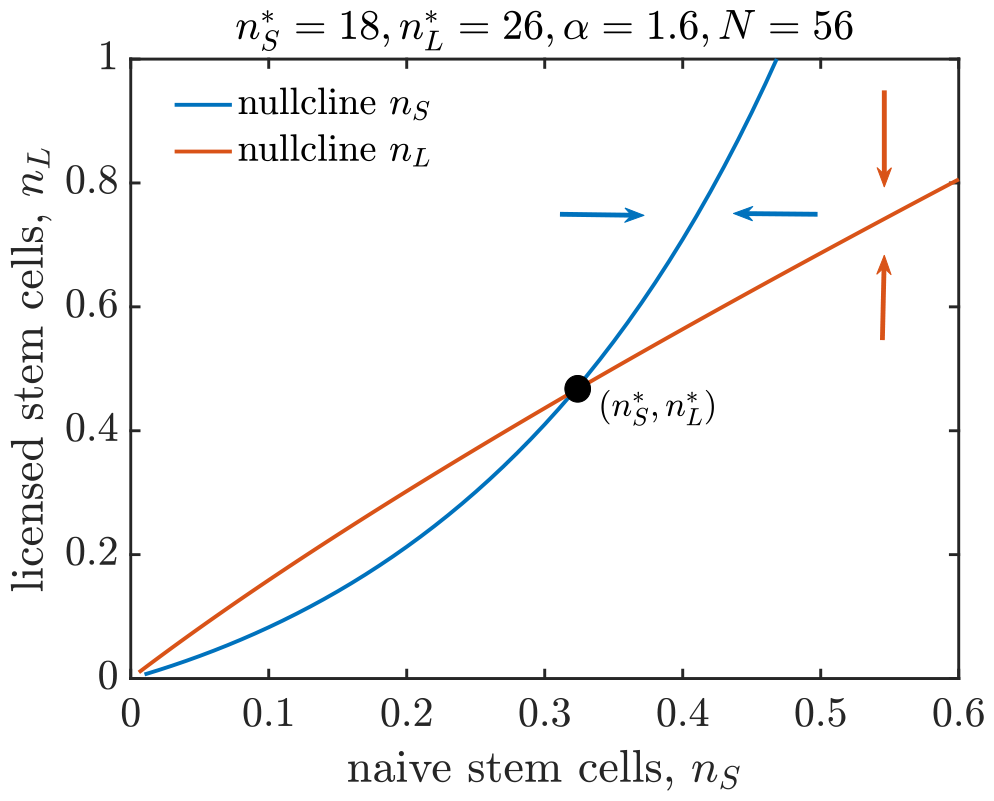
Phase portrait of the v*SL* model. The steady states ((0, 0) and 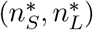) are the intersection points of the nullclines (zero-growth curves) of *n*_*S*_ (blue) and *n*_*L*_ (orange). Arrows represent the directions of growth outside the nullclines, displaying 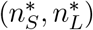 as a stable steady state.

Let **f** (*n*_*S*,_ *n*_*L*_), [*f*_1_(*n*_*S*,_ *n*_*L*_), *f*_2_ (*n*_*S*,_ *n*_*L*_)] with 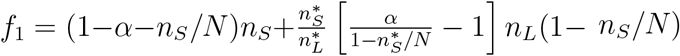 and 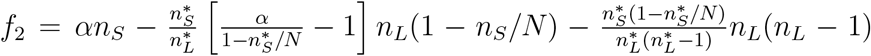.Defining *ϕ*_*S*_ *= n*_*S*_ /*N and ϕ*_*L*_ *= n*_*L*_ /*N*, the Jacobian of **f** at the steady state is given by

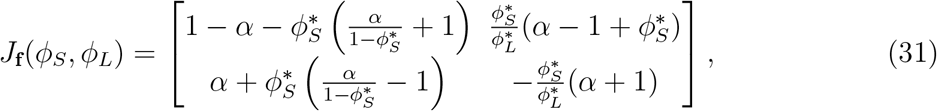

where we have approximated 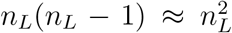. Eigenvalue calculation yields two real, negative eigenvalues for *α >* 1, which proves the linear stability of the nontrivial steady state. The stability of the steady state also becomes evident upon inspection of the phase portrait of the v*SL* model in Fig. 9.

#### 2. Stochastic implementation of the vSL model

We follow a similar procedure than for the *SL* model in Sec. VII A 2, with the constraint that for the v*SL* model analytical solutions of the Lyapunov equation (Eq. (19)) are rather cumbersome. Instead, we solve the Lyapunov equation numerically for each parameter set to find the second moments of the distributions of stem cell numbers in quasi-steady state. The second moments of the distribution in quasi-steady state obtained via the LNA are in good agreement with SSA calculations (Fig. 10).

**FIG. 10.**
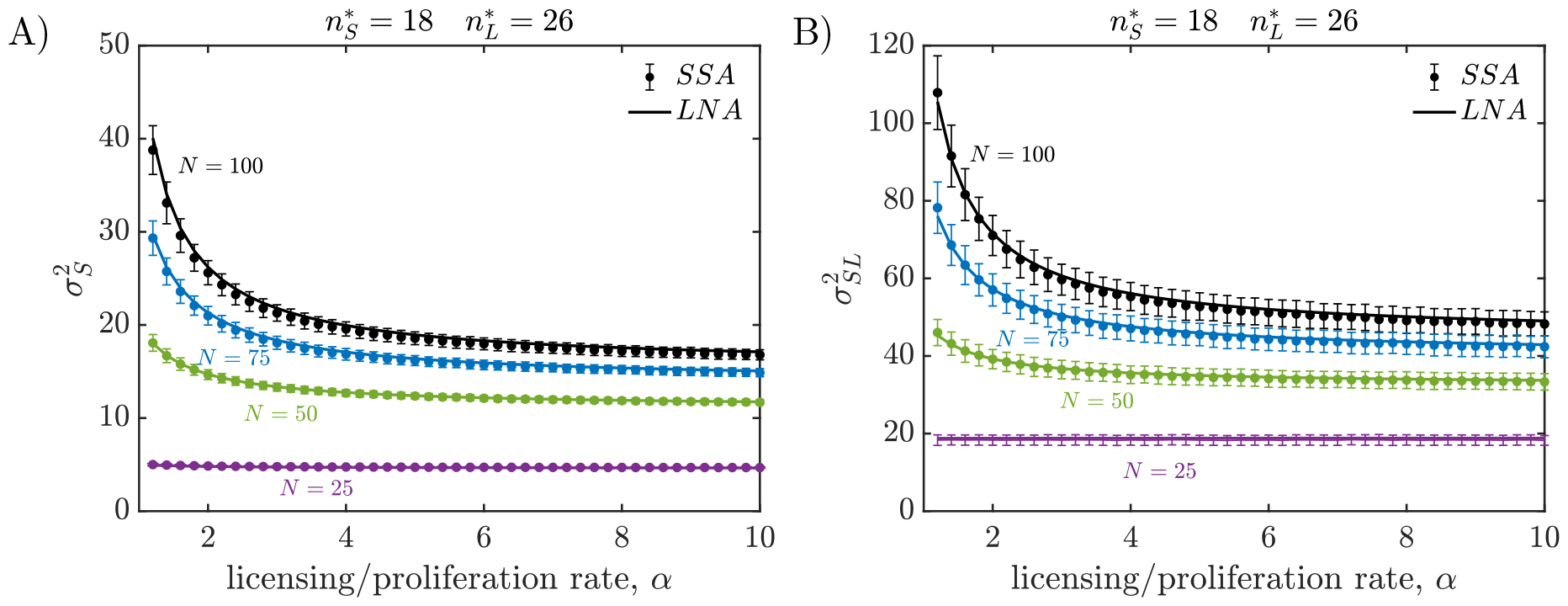
Second moments of the distribution of naive. (**A**) and total (**B**) stem cell numbers in quasi-steady state, as a function of *α* for the *υ SL* model with different carrying capacities *N*. The variance measurements (dots) are averaged over 1 *×* 10^3^ realisations of the SSA with 1 *×* 10^4^ time points. Error bars show one standard deviation in the ensemble of realisations. The LNA predictions (lines), obtained from numerically solving the Lyapunov equation (Eq. (19)), are in agreement with the SSA measurements.

## References

[1] E. Laurenti and B. Göttgens, From haematopoietic stem cells to complex differentiation landscapes, Nature 553, 418 (2018).

[2] K. A. Moore and I. R. Lemischka, Stem cells and their niches, Science 311, 1880 (2006).

[3] R. García-Tejera, L. Schumacher, and R. Grima, Regulation of stem cell dynamics through volume exclusion, Proceedings of the Royal Society A 478, 20220376 (2022).

[4] B. Corominas-Murtra, C. L. Scheele, K. Kishi, S. I. Ellenbroek, B. D. Simons, J. Van Rheenen, and E. Hannezo, Stem cell lineage survival as a noisy competition for niche access, Proceedings of the National Academy of Sciences 117, 16969 (2020).

[5] C. Parigini and P. Greulich, Homeostatic regulation of renewing tissue cell populations via crowding control, arXiv preprint 2301.05321 (2023).

[6] Y. Kitadate, D. J. Jörg, M. Tokue, A. Maruyama, R. Ichikawa, S. Tsuchiya, E. Segi-Nishida, T. Nakagawa, A. Uchida, C. Kimura-Yoshida, et al., Competition for mitogens regulates spermatogenic stem cell homeostasis in an open niche, Cell Stem Cell 24, 79 (2019).

[7] S. Yoshida, Open niche regulation of mouse spermatogenic stem cells, Development, growth & differentiation 60, 542 (2018).

[8] D. J. Jörg, Y. Kitadate, S. Yoshida, and B. D. Simons, Stem cell populations as self-renewing many-particle systems, Annual Review of Condensed Matter Physics 12, 135 (2021).

[9] R. R. Stine and E. L. Matunis, Stem cell competition: finding balance in the niche, Trends in cell biology 23, 357 (2013).

[10] E. Hannezo, A. Coucke, and J.-F. Joanny, Interplay of migratory and division forces as a generic mechanism for stem cell patterns, Physical Review E 93, 022405 (2016).

[11] K. H. Vining and D. J. Mooney, Mechanical forces direct stem cell behaviour in development and regeneration, Nature reviews Molecular cell biology 18, 728 (2017).

[12] Y. Zhang, H. Wei, and W. Wen, Phase separation and mechanical forces in regulating asymmetric cell division of neural stem cells, International Journal of Molecular Sciences 22, 10267 (2021).

[13] M. Renardy, A. Jilkine, L. Shahriyari, and C.-S. Chou, Control of cell fraction and population recovery during tissue regeneration in stem cell lineages, Journal of theoretical biology 445, 33 (2018).

[14] A. D. Lander, K. K. Gokoffski, F. Y. M. Wan, Q. Nie, and A. L. Calof, Cell lineages and the logic of proliferative control, PLoS biology 7, e1000015 (2009).

[15] I. A. Rodriguez-Brenes, D. Wodarz, and N. L. Komarova, Stem cell control, oscillations, and tissue regeneration in spatial and non-spatial models, Frontiers in oncology 3, 82 (2013).

[16] M. Amoyel, K.-H. Hillion, S. R. Margolis, and E. A. Bach, Somatic stem cell differentiation is regulated by pi3k/tor signaling in response to local cues, Development 143, 3914 (2016).

[17] A. C. Yuen, K.-H. Hillion, R. Wang, and M. Amoyel, Germ cells commit somatic stem cells to differentiation following priming by pi3k/tor activity in the drosophila testis, PLoS genetics 17, e1009609 (2021).

[18] L. Weinberger, M. Ayyash, N. Novershtern, and J. H. Hanna, Dynamic stem cell states: naive to primed pluripotency in rodents and humans, Nature reviews Molecular cell biology 17, 155 (2016).

[19] J. S. Haug, X. C. He, J. C. Grindley, J. P. Wunderlich, K. Gaudenz, J. T. Ross, A. Paulson, K. P. Wagner, Y. Xie, R. Zhu, et al., N-cadherin expression level distinguishes reserved versus primed states of hematopoietic stem cells, Cell stem cell 2, 367 (2008).

[20] T. Nakagawa, D. J. Jörg, H. Watanabe, S. Mizuno, S. Han, T. Ikeda, Y. Omatsu, K. Nishimura, M. Fujita, S. Takahashi, et al., A multistate stem cell dynamics maintains homeostasis in mouse spermatogenesis, Cell Reports 37, 109875 (2021).

[21] F. d. S. e Melo and F. J. de Sauvage, Cellular plasticity in intestinal homeostasis and disease, Cell Stem Cell 24, 54 (2019).

[22] M. A. Goodell, H. Nguyen, and N. Shroyer, Somatic stem cell heterogeneity: diversity in the blood, skin and intestinal stem cell compartments, Nature reviews Molecular cell biology 16, 299 (2015).

[23] J. Nichols and A. Smith, Naive and primed pluripotent states, Cell stem cell 4, 487 (2009).

[24] Q.-L. Ying, J. Wray, J. Nichols, L. Batlle-Morera, B. Doble, J. Woodgett, P. Cohen, and A. Smith, The ground state of embryonic stem cell self-renewal, nature 453, 519 (2008).

[25] K. Takahashi and S. Yamanaka, A decade of transcription factor-mediated reprogramming to pluripotency, Nature reviews Molecular cell biology 17, 183 (2016).

[26] L. J. Greenspan, M. De Cuevas, and E. Matunis, Genetics of gonadal stem cell renewal, Annual review of cell and developmental biology 31, 291 (2015).

[27] K. F. Lenhart and S. DiNardo, Somatic cell encystment promotes abscission in germline stem cells following a regulated block in cytokinesis, Developmental cell 34, 192 (2015).

[28] J. L. Leatherman and S. DiNardo, Zfh-1 controls somatic stem cell self-renewal in the drosophila testis and nonautonomously influences germline stem cell self-renewal, Cell stem cell 3, 44 (2008).

[29] J. J. Fabrizio, M. Boyle, and S. DiNardo, A somatic role for eyes absent (eya) and sine oculis (so) in drosophila spermatocyte development, Developmental biology 258, 117 (2003).

[30] M. Amoyel, B. D. Simons, and E. A. Bach, Neutral competition of stem cells is skewed by proliferative changes downstream of hh and hpo, The EMBO journal 33, 2295 (2014).

[31] M. Issigonis, N. Tulina, M. De Cuevas, C. Brawley, L. Sandler, and E. Matunis, Jak-stat signal inhibition regulates competition in the drosophila testis stem cell niche, Science 326, 153 (2009).

[32] S. R. Singh, Z. Zheng, H. Wang, S.-W. Oh, X. Chen, and S. X. Hou, Competitiveness for the niche and mutual dependence of the germline and somatic stem cells in the drosophila testis are regulated by the jak/stat signaling, Journal of cellular physiology 223, 500 (2010).

[33] J. E. Till, E. A. McCulloch, and L. Siminovitch, A stochastic model of stem cell proliferation, based on the growth of spleen colony-forming cells, Proceedings of the National Academy of Sciences of the United States of America 51, 29 (1964).

[34] A. M. Klein and B. D. Simons, Universal patterns of stem cell fate in cycling adult tissues, Development 138, 3103 (2011).

[35] N. A. Robertson, E. Latorre-Crespo, M. Terrada-Terradas, A. C. Purcell, B. J. Livesey, J. A. Marsh, L. Murphy, A. Fawkes, L. MacGillvray, M. Copland, et al., Longitudinal dynamics of clonal hematopoiesis identifies gene-specific fitness effects, bioRxiv (2021).

[36] C. Parigini and P. Greulich, Universality of clonal dynamics poses fundamental limits to identify stem cell self-renewal strategies, Elife 9, e56532 (2020).

[37] S. Rulands, F. Lescroart, S. Chabab, C. J. Hindley, N. Prior, M. K. Sznurkowska, M. Huch, A. Philpott, C. Blanpain, and B. D. Simons, Universality of clone dynamics during tissue development, Nature physics 14, 469 (2018).

[38] P. Greulich and B. D. Simons, Dynamic heterogeneity as a strategy of stem cell self-renewal, Proceedings of the National Academy of Sciences 113, 7509 (2016).

[39] S. Hasan, P. Hétié, and E. L. Matunis, Niche signaling promotes stem cell survival in the drosophila testis via the jak–stat target diap1, Developmental biology 404, 27 (2015).

[40] D. T. Gillespie, A general method for numerically simulating the stochastic time evolution of coupled chemical reactions, Journal of computational physics 22, 403 (1976).

[41] D. Schnoerr, G. Sanguinetti, and R. Grima, Comparison of different moment-closure approx-imations for stochastic chemical kinetics, The Journal of Chemical Physics 143, 11B610 1 (2015).

[42] M. Assaf and B. Meerson, Extinction of metastable stochastic populations, Physical Review E 81, 021116 (2010).

[43] M. Ziehm, M. D. Piper, and J. M. Thornton, Analysing variation in d rosophila aging across independent experimental studies: a meta-analysis of survival data, Aging Cell 12, 917 (2013).

[44] M. Ziehm and J. M. Thornton, Unlocking the potential of survival data for model organisms through a new database and online analysis platform: S urv c urv, Aging cell 12, 910 (2013).

[45] A. J. McKane and T. J. Newman, Stochastic models in population biology and their deter-ministic analogs, Physical Review E 70, 041902 (2004).

[46] C. Cianci, S. Smith, and R. Grima, Capturing brownian dynamics with an on-lattice model of hard-sphere diffusion, Physical Review E 95, 052118 (2017).

[47] S. Yoshida, Heterogeneous, dynamic, and stochastic nature of mammalian spermatogenic stem cells, Current topics in developmental biology 135, 245 (2019).

[48] N. G. Van Kampen, Stochastic processes in physics and chemistry, Vol. 1 (Elsevier, 1992).

[49] D. Schnoerr, G. Sanguinetti, and R. Grima, Approximation and inference methods for stochastic biochemical kinetics—a tutorial review, Journal of Physics A: Mathematical and Theoretical 50, 093001 (2017).

[50] J. Elf and M. Ehrenberg, Fast evaluation of fluctuations in biochemical networks with the linear noise approximation, Genome research 13, 2475 (2003).

